# Target Detection Modulates EEG Spectral Correlates of Memory Encoding

**DOI:** 10.1101/2023.04.24.538090

**Authors:** Adam W. Broitman, Khena M. Swallow

## Abstract

The current study investigates whether changes in scalp electroencephalographic (EEG) activity over time reflect the effects of target detection and divided attention on memory encoding. We recorded EEG activity in 61 young adults as they memorized lists of words either under full attention (single-task) or while performing a secondary task (dual-task). In both cases, colored squares appeared with each word. However, in the dual-task condition participants also pressed a button when the colored squares were in a predefined color (target) but made no response when the squares were in a different color (distractor). Subsequent memory effects in the alpha (8-12 Hz) and high gamma (50-100 Hz) frequency bands changed throughout the trial, and these effects differed across conditions. Prior to word presentation, high gamma activity was associated with encoding success in the target and single-task conditions, but not in the distractor conditions. In contrast, alpha band activity decreased following word presentation, and these decreases were greater for successfully encoded words in the target condition than in the distractor or single-task conditions. The results are consistent with the view that alpha and gamma activity reflect distinct neural processes which both contribute to memory formation, but are differentially sensitive to task demands and momentary shifts in attention.

## Introduction

Most everyday tasks require changes in attention over spatial locations or item features. However, attention can also vary across time (Rohenkohl et al., 2011), and this can modulate memory encoding in a number of ways. For example, changes in task demands, such as when a person performs multiple tasks simultaneously, can impact cognitive load and interfere with the encoding of new memories (Craik et al., 1996). Similarly, phasic changes in attention (e.g., due to the detection of a salient or important stimulus) can dramatically influence memory encoding from one moment to the next (Mather et al., 2016; Swallow & Jiang, 2010). Spectral analyses of electroencephalogram (EEG) recordings have provided critical insights into the influence of attention on memory processes, linking specific changes in neural activity with memory formation (e.g., Klimesch et al., 1998; Long, Burke, & Kahana, 2014). However, there remain several key questions about how EEG activity reflects attention-memory interactions. Specifically, it is unclear whether the neural activity associated with encoding similarly predicts memory formation under divided attention, or while individuals respond to behaviorally relevant stimuli. Furthermore, it is unclear whether any potential effects of attention manifest throughout the encoding period, or only following the presentation of task-related stimuli. The current study investigates these questions using spectral decomposition of EEG in a study of free recall.

### The effects of attention and successful encoding on EEG activity

Attention can influence episodic encoding at multiple time scales, with some effects constrained to brief intervals (Swallow & Jiang, 2010; Swallow & Jiang, 2011; Makovski et al., 2011) and others reflecting longer lasting attentional states and task demands (e.g., Troyer & Craik, 2000). Investigations of the effects of temporal variations in attention on encoding are aided by methods with a high degree of temporal resolution, such as scalp electroencephalography (EEG). Consequently, EEG has been extensively used to study the mechanisms involved in attentional orienting and successful episodic memory encoding. These studies suggest that characterizing activity in alpha (8-12 Hz) and gamma (30+ Hz) frequency bands could provide novel insights into how dual-task demands and target detection modulate encoding.

Prior EEG studies have identified patterns of spontaneous brain activity, called neural *subsequent memory effects* (SMEs), which can help predict whether items presented in that moment will be remembered or forgotten (Paller et al., 1987; Karis et al., 1984; Fabiani et al., 1986; 1990). Neural SMEs typically consist of a complex combination of features that may reflect multiple neurocognitive mechanisms (Kilner et al., 2005; Voytek & Knight, 2015), and could be sensitive to various aspects of attention.

EEG studies of free recall have reported that decreases in power spectral density (PSD) within lower frequencies (e.g., alpha, 8-12 Hz), combined with increases in power in higher frequencies (e.g., gamma, 30+ Hz), are associated with successful encoding (Long, Burke, & Kahana, 2014; Sederberg et al., 2003). However, high and low frequency bands may reflect distinct neural processes that are differentially sensitive to a variety of task demands. For example, Sederberg et al. (2006) reported that in free recall experiments gamma SMEs weaken across serial positions as subjects process multiple items from early list positions, but alpha SMEs become stronger. In an experiment where participants encoded words under full attention, or while performing a secondary semantic decision task on the to-be-encoded word (e.g., “is this item living/nonliving”), Broitman et al. (*under review*) found that gamma activity was only associated with encoding success when subjects encoded words under full attention, while alpha SMEs were observed under both dual-task and full attention conditions. Using data from the same experiment, Long & Kahana (2017) found that gamma predicted the tendency to temporally cluster recalls when words were encoded under full attention, but not when subjects performed the secondary judgment task. Finally, there is some evidence that gamma-band SMEs emerge prior to or immediately following trial onset (-300 to 500 ms), while alpha band SMEs occurred later in the trial (1000 to 1500 ms; Noh et al., 2014; Long et al., 2014, but see Weidemann et al., 2021), suggesting that they reflect different aspects of task performance.

A potential explanation for these findings is that gamma and alpha activity are modulated by different components of attention that influence memory encoding: the cognitive system’s level of task engagement and the orienting to and processing of a stimulus once it appears. Investigations into how alpha and gamma power change during attention tasks support this possibility. Alpha suppression occurs in scalp sites that are contralateral to the attended visual field (Sauseng, et al., 2005) and occurs broadly across the scalp following the presentation of targets in a detection task (Klimesch et al., 1998; de Graaf & Thompson, 2020; He et al., 2021). Further, alpha suppression is associated with pupil dilations (Compton et al. 2021), which may reflect the LC system’s involvement in disrupting alpha oscillations throughout the brain (Dahl et al., 2022). Thus, alpha power should be modulated by orienting toward behaviorally relevant stimuli, like targets, after they appear. The effect could be particularly strong when encoding is successful.

Like activity in the alpha band, gamma band activity may reflect the task-oriented deployment of attention. However, in EEG, gamma activity can capture changes in attentional states over a longer time frame. In addition to studies showing that gamma power decreases during list encoding (Sederberg et al., 2006), other studies have observed gamma power gains in response to increasing sensory processing demands (Von Stein & Sarnthein, 2000), as well as cognitive task difficulty (Rietschel et al., 2012). In a study combining fMRI and iEEG measures, Kucyi et al. (2020) found that high gamma oscillations occurred earlier in the dorsal attention network when targets were accurately detected. This may suggest a role of gamma band activity in the goal-directed establishment of cognitive states needed for task performance (Clayton, et al., 2015). If this is the case, then gamma band activity may be sensitive to longer lasting increases in attentional demands from divided attention and may predict how brief increases in attentional demands from target detection impact encoding success.

### The Attentional Boost Effect

The combined effects of attentional states on memory encoding, specifically as they relate to cognitive load and orienting, are present in the *attentional boost effect* (ABE; Swallow & Jiang, 2010). In the standard attentional boost effect task, participants study a continuous stream of items (e.g., faces, words) while simultaneously monitoring an unrelated stream of coinciding stimuli (e.g., a pair of squares that appear simultaneously with the item). Participants press a button when the squares are in a predefined *target* color rather than a *distractor* color (e.g., a pair of blue rather than red squares). Despite increased demands on attention from the target (e.g., Duncan, 1980), memory for an item that coincides with the target is enhanced relative to memory for an item that appears with a distractor or on its own (Mulligan, et a., 2014; Swallow & Jiang, 2014). Importantly, the task requires participants to sustain attention over time, quickly respond to behaviorally relevant stimuli, and divide attention across two stimulus streams and tasks. This makes it an ideal paradigm to study how different sources of variability in attention over time can influence memory encoding and its associated neural signatures.

Although the attentional boost effect has been extensively replicated under varying experimental conditions (see, Swallow, Broitman, Riley & Turker, 2022 for a review), the specific attentional mechanisms that are modulated with this effect remain unclear. Accounts for the attentional boost effect often focus on the effects of target detection on how well other information is processed. For example, the Dual-Task Interaction model (DTI; Swallow & Jiang, 2013) proposes that the detection of a behaviorally relevant stimulus briefly activates the Locus Coeruleus (LC) system, which increases the brain’s sensitivity to external stimuli and boosts episodic encoding (Takeuchi et al., 2016) in a process known as *temporal selection*. Other work similarly suggests that the attentional boost effect arises from enhancements to early components of episodic encoding following target detection (Mulligan & Spataro, 2015). Another possibility is that detecting a target could briefly alleviate the cognitive demands of performing two tasks simultaneously by reducing conflict between them (e.g., by increasing processing efficiency). If such a state occurs with target detection, then increases in processing efficiency might allow people to encode background items in a manner similar to that of full attention. These competing frameworks provide mechanistic explanations of attentional boost effect, but speak little to the potential role of preexisting attentional states (i.e., proactively sustained attention to the task) on target-related memory enhancements.

### The current study

The current experiment aims to investigate the effects of target detection on SMEs associated with attentional orienting and maintaining task readiness under divided attention conditions. We recorded scalp EEG while participants performed a delayed free recall task, with some participants encoding wordlists under full attention while others additionally performed a visual target detection task during encoding. Consistent with previous literature, we expected target detection under dual-task conditions to enhance subsequent recall of the concurrently presented words (e.g., Broitman & Swallow, 2023; Mulligan, et al. 2014). Of greater interest, however, was the impact of these manipulations on EEG power spectral density in the alpha and gamma frequency bands at different periods of the trial.

First, we expected single-task conditions to replicate the previously reported SMEs of decreased alpha activity coupled with increased gamma power. However, these patterns may not hold when participants encode words while performing a second, target detection task. Prior studies have demonstrated that low frequency (alpha band) power is sensitive to external attentional orienting (Klimesch et al., 1998; de Graaf & Thompson, 2020; He et al., 2021), suggesting that alpha SMEs will be modulated by target detection. Further, because previous work suggests that cognitive load decreases gamma SMEs (Long & Kahana, 2017; Broitman et al., *Under Review*), gamma SMEs may be modulated by both divided attention and target detection. Importantly, because prior work utilized secondary tasks that directly altered how the words were processed, this would be the first demonstration that gamma band SMEs are sensitive to the distribution of attention over multiple tasks and stimuli.

Of critical interest was at what time points our attentional manipulations influenced the spectral SMEs. Given the relationship between alpha oscillations and the locus coeruleus system (Compton et al., 2021), our hypothesis is that target detection will manifest in alpha power suppression *following* targets, and that greater alpha suppression will be associated with encoding success (see Klimesch et al., 1998; de Graaf & Thompson, 2020). Alternatively, if target detection alleviates the demands of performing two tasks simultaneously, then SMEs on target trials may begin to resemble those observed under single-task conditions, but only *after* a target occurs. An additional possibility is that encoding success on target- and distractor-trials depends on one’s cognitive state, as indicated by power spectral density SMEs, *prior to stimulus onset.* Such a finding would suggest that preexisting mental states modulate the effects of target detection on memory encoding. To investigate these possibilities, we compared EEG spectral activity recorded both before and after stimulus onset during single vs. dual task trials, as well as target vs. distractor trials within the dual task condition.

## Method

### Participants

Right-handed, neurologically healthy participants aged 18-40 were recruited from the Cornell campus and greater Ithaca community. Participants were compensated with either $10/per hour of participation or psychology course credit. All participants had normal or corrected-to-normal vision, and normal color vision was verified with the Hardy, Rand & Ritler pseudoisochromatic color blindness test (Richmond Products, Albuquerque, NM, USA). The experimental procedures were reviewed and approved by Cornell’s Institutional Review Board. All participants provided informed consent and were debriefed about the purpose of the experiment upon completion.

To ensure the replicability of the SMEs and estimate an effect size, we first collected EEG data from 12 participants in a single-task free recall experiment. Based on these data, a target sample size of 40 participants in the dual-task experiment was selected *a priori* using G*Power 4.1 (Faul, Erdfelder, Lang, & Buchner, 2007), assuming an effect of size *d* = 0.59 with a power (1 − ß) of .95 in a two-tailed paired samples *t*-test. Data from 1 participant whose performance did not meet the inclusion criteria in the detection task (>80% hits, <10% false alarms) or recall task (>100, or 22%, words correctly recalled) were replaced, resulting in a final sample size of 40 in the dual-task condition. Following the collection of data from the 40 dual-task participants, a final single-task target sample size of 20 was selected to approximately match the number of trials in each dual-task cue condition. Data from 3 participants who failed to meet the inclusion criteria of at least 100 correctly recalled words (23% of all words) were excluded and replaced, resulting in a final sample size of 21.

### Experiment Equipment and Materials

Participants sat unconstrained in a normally lit interior room, approximately 90 cm away from a View-Sonic E70fB 17′′ CRT monitor (1024×768 pixels, 75Hz refresh rate) controlled by a Dell PC and responded on a keyboard. All experiments were programmed in PsychoPy (Pierce et al., 2019).

Word lists were created using methods derived from the Penn Electrophysiology of Encoding and Retrieval Study (see Broitman, Kahana, & Healey, 2020; Kahana et al., *in press*). Words were chosen from a list of 1638 nouns, and lists were constructed such that adjacent words would have varying degrees of semantic similarity with one another. Inter-item similarity was constructed using the Word Association Space (WAS; Steyvers, Shiffrin, & Nelson, 2005). Each list included two word pairs chosen from each of five similarity bins ranging from low- to high-similarity, with the words from one pair placed at adjacent serial positions and the words from the other pair separated by at least two serial positions (see Kahana et al., *in press*). Because this method produced a list length of 20 words, the lists were then truncated to a length of either 18 (single-task condition) or 16 (dual-task condition).

Prior to data collection, participants were fitted with a 64-Channel Biosemi ActiveTwo geodesic cap. Scalp EEG data was collected at a sampling rate of 512 Hz. Participants were instructed to minimize blinks during stimulus presentation, and to limit muscle movements when performing the task.

### Procedure

#### Single-Task Encoding Condition

Participants completed 24 cycles of the continuous encoding task, with each cycle followed by a math distractor task, and a free recall task. Each cycle of the continuous encoding task used a different list of words. The lists each consisted of 18 words, resulting in a total of 432 words across the 24 lists. Each of the 28 wordlist presentations began with a fixation presented for 2000 ms at the center of the screen. Each trial (duration = 1250-1750 ms) consisted of a word presented at the center of the screen with a square above and below it for 100 ms. Both squares presented were either red (RGB: 255, 0, 0) or green (RGB: 0, 255, 0). Participants were told to ignore the colored squares and focus only on encoding the words. Following the presentation of the word and squares, the word appeared by itself onscreen for an additional 400 ms, followed by a jittered (i.e., variable) interstimulus interval of 750-1250 ms during which the screen was blank. Red and green squares appeared at each serial position with equal frequency across wordlists, and squares of the same color were never presented on more than four consecutive trials within a wordlist.

Following the presentation of the words, participants performed simple arithmetic problems (e.g., -1 + 3 + 2 = ?) for 30 s. For this task, participants were instructed to correctly solve as many math problems as they could during the 30 s period. Following the math distractor period, participants were instructed to freely recall and type out as many words from the previous list as they could remember for a period of 75 s. For the recall period, participants used their keyboards to type out each word that they remembered from the immediately preceding list. Participants pressed the RETURN key after entering a word.

#### Dual-Task Encoding Condition

The procedure for the dual-task condition was identical to that of the single-task condition except for the following details. Because the attentional boost effect paradigm requires memory performance to be sufficiently high to detect an effect of target detection on encoding (Broitman & Swallow, 2020; Hutmacher & Kuhbandner, 2020), and because performing a secondary task at encoding can impair free recall performance (Craik, 1996; Fernandez & Moskovitch, 2000), we reduced the number of items per list from 18 to 16. To better match the single- and dual-task conditions on the overall number of trials, we also increased the number of word lists from 24 to 28, resulting in a total of 448 trials per subject. As with the single-task condition, each word was presented with a set of red or green squares. However, instead of being instructed to ignore the squares and focus only on encoding the words, participants were instructed to press the spacebar if the squares were a predefined target color. Half of the participants were instructed to press the spacebar if the squares were green, while the other half were instructed to press the spacebar if the squares were red. Participants had a 1000 ms deadline to respond to targets. Because the attentional boost effect diminishes once trial durations exceed 2000 ms (Mulligan et al, 2014), trial durations in both experimental conditions were limited to 1750 ms in order to optimize the effects of target detection on memory encoding.

### Data Analysis

#### EEG preprocessing

EEG preprocessing and analysis was performed using the MNE software package (Gramfort et al., 2013) in conjunction with custom python scripts. To remove electrical noise that may have been present in the data, a bandpass filter was applied between 0.1 and 100 Hz, and a notch filter was applied at 60 Hz. The filtered data was then visually inspected to identify any electrodes displaying erratic activity.

On average, fewer than one channel per participant was interpolated and excluded. Excessively noisy channels were identified and interpolated using MNE’s spherical spline interpolation method. Next, EEG artifacts consistent with eye and muscle movements were detected and removed with independent component analysis (ICA) on the artifact-corrected data using a “fast ICA” algorithm (Ablin et al., 2018). Components were selected by visually inspecting their topography (e.g, concentrated near the eyes) and by comparing component activations with the raw data feed to determine if a component was temporally aligned with a stereotyped artifact (e.g., blinks or head movements). Approximately 3 ICA components were removed per participant. Finally, we identified bad epochs as those with an extremely high variance (|*z|* > 3), and excluded them from further analysis.

#### EEG spectral density analyses

Following EEG preprocessing, we segmented the EEG data into epochs beginning 500 ms prior to stimulus onset and ending 1000 ms post-stimulus onset. For each electrode, PSD on each trial was computed at 18 logarithmically spaced frequencies ranging from 2-100 Hz using a Morlet wavelet transformation with a wavelet number of five. Power values were then log-transformed and decimated by a factor of 32, resulting in a final sampling rate of 16 Hz. This produced PSD values for 25 time points between -500 and 1000 ms relative to stimulus onset at each trial. Power within each channel and frequency was then Z-scored using the mean and standard deviation of power values across data points from all trials completed by the participant. This method of z-scoring has the advantage of removing variance in mean power across scalp sites and permits comparison of pre-stimulus periods (which are sometimes used for baseline correction) across conditions. However, the topographies of alpha and gamma band SMEs are typically similar across a variety of normalization methods (Weidemann & Kahana, 2021; Hanslmayr, Spitzer, & Bauml, 2009; Fellner, Bauml, & Hanslmayr, 2013). Power values were subsequently averaged within five frequency bands: Theta (4-8 Hz), Alpha (8-12 Hz), Beta (12-30 Hz), Low Gamma (30-50 Hz), and High Gamma (50-100 Hz). For analyses of trial-level power, the PSD estimates were averaged across all time points in the trial. For time series analyses, the PSD estimates were averaged over 500 ms time bins (*pre-trial*, -500-0 ms; *early trial*, 0-500 ms; *late trial*, 500-1000 ms).

#### Statistical analysis

Two-tailed paired-samples t-tests were conducted to check for the presence of previously characterized subsequent memory effects. Further analyses relied on linear mixed effects models, and contrast analyses using the lmer function (lme4; Bates, et al., 2015) and the emmeans (Russell, 2018) package in R.

Two-tailed t-tests were also used to compare free recall rates between the trial conditions. Additional features of free recall behavior were investigated using linear mixed effects models with fixed effects of task condition (single vs. dual), serial position, and cue condition (target vs. distractor). All models included random intercepts for participant, and were further characterized using the emmeans package.

To better understand whether spectral activity was differentiated by trial type, subsequent memory, or time within a trial, we constructed two linear mixed regression models with either alpha or high gamma power (PSD as the dependent variable. Each model included fixed effects for subsequent memory (remembered/forgotten), trial type (target/distractor/single task), and time bin (pretrial/early trial/late trial). Random intercepts were included for participant by subsequent memory, time, and trial type, using the following notation:

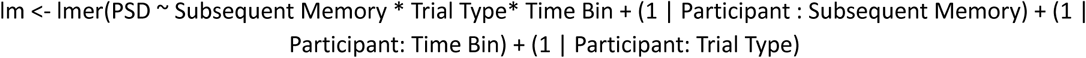

The models will henceforth be referred to as the *Alpha MLM* and the *Gamma MLM*. For all analyses with these models, specific comparisons (e.g., between target and distractor conditions) were investigated using custom contrasts in emmeans. Comparisons of single- to dual-task were performed using a contrast that averaged over the target and distractor trials for the dual-task condition (i.e., target: 0.5; distractor: 0.5; single-task: -1). Interactions were investigated using a standard difference of differences approach (e.g., for the interaction of cue type and subsequent memory: remembered target: 1; forgotten target: -1; remembered distractor: -1; forgotten distractor: 1).

## Results

### Behavioral Results

#### Detection Task Performance

Among participants in the dual-task encoding condition, detection task performance was high, with a hit rate of 98.7% (SD = 1.5%) and a false alarm rate of 3% (SD = 1.9%). Target response times averaged 515 ms (SD = 82 ms), and did not differ depending on whether the coinciding word was subsequently recalled or not, *t*(40) = .01, p = .98.

#### Free Recall Performance

In Figure 1c, recall rates are plotted separately for target, distractor, and single-task encoding conditions in each serial position. In the dual-task condition, a two-tailed paired samples t-test indicated that target-paired words were recalled more frequently than words presented with distractors, *t*(39) = 4.2, *p* < .001, *d* = .68. Participants also were more likely to initiate recall with a target-paired item. The probability of first recall was significantly higher for target-paired words, even after controlling for the attentional boost effect within each participant, *F*(1, 76) = 53.4, *p* < .0001, η^2^ = .41. These data indicate that words presented with targets during encoding were more accessible during retrieval and replicate prior findings (Broitman & Swallow, 2023).

**Figure 1.**
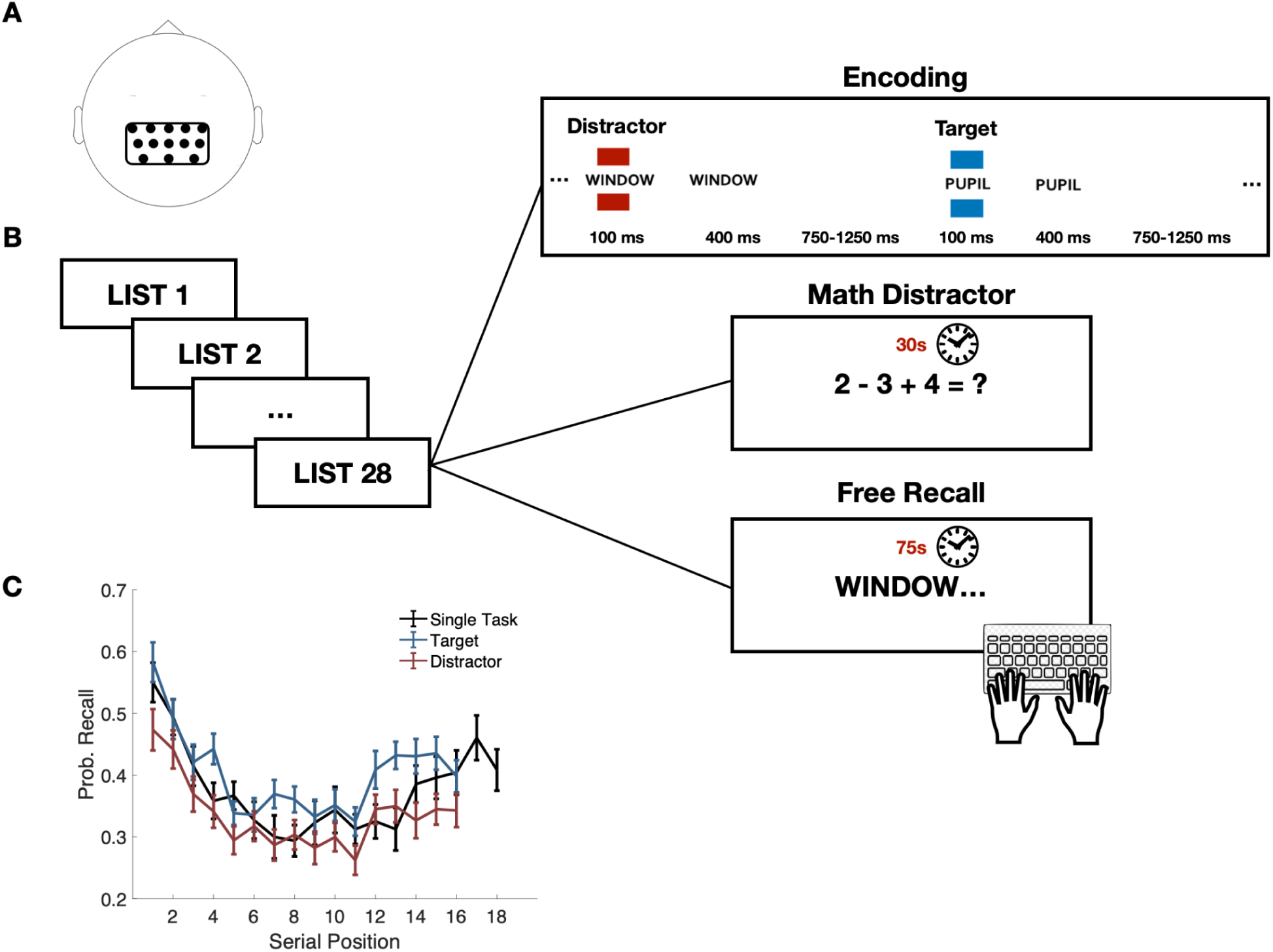
**A)** Bilateral parietal ROI selected for analysis. **B)** Illustration of the experimental task. Participants were presented with word lists and colored squares, and instructed to either focus only on the words (single-task) or to perform target detection on the colored squares while encoding the words (dual-task). Participants in the dual-task condition encoded 28 lists of 16 words, and participants in the single-task condition encoded 24 lists of 18 words. Each word list presentation was followed by a 30 s math distractor test and a 75 s retrieval period. Color cues are depicted in red and blue for consistency with the figures, though they were red and green in the actual experiment. **C)** Free recall performance for words presented at each serial position in the single- and dual-task encoding conditions, with the latter separated into target and distractor cue conditions. Error bars reflect the standard error of the mean across participants.

We did not detect any serial positioning or temporal contiguity effects that could account for differences in EEG activity between target and distractor cue conditions. A mixed linear regression with recall rates as the dependent variable revealed no interaction between cue condition and serial position, *F*(1,1209) = .60, *p* = .86, η^2^ < .01. The target-related memory advantage was therefore approximately uniform across serial positions. Further, no differences between target and distractor paired words were observed in two metrics that capture temporal clustering dynamics in recall: the temporal clustering score and forward transition asymmetry. The temporal clustering score (Polyn et al., 2009) reflects the proportion of lags to the available recall transitions (i.e., to any not-yet-recalled items from the study list) that the actual transition lag is *less than*. Forward transition asymmetry (FTA) indicates the propensity to successively retrieve adjacent list items in the forward, rather than backward, direction (Howard & Kahana, 2002). Two-tailed paired-samples t-tests demonstrated that temporal clustering scores were similar among transitions from target- and-distractor cue conditions, t(38) = .58, p = .56, d = .11, as were estimates of FTA, t(38) = 1.2, p = .25, d = .22. These results are consistent with those reported in Broitman & Swallow (2023), in which target detection did not interact with temporal contiguity or serial position effects in recall.

Among participants in the single-task encoding condition, recall rates did not significantly differ from the overall dual-task recall rates, *t*(59) = .40, *p* = .68, *d* = .09, or from the separate target or distractor cue conditions, *t*(59)’s <= 1.6, *p* > .12. List lengths in the dual-task condition were shortened to ensure recall was sufficiently high in this condition, which may explain the lack of a difference from single-task recall rates. However, we observed greater levels of temporal clustering among participants in the single-task condition than those in the dual-task condition, t(59) = 2.5, p = .02, d = .70. This result is consistent with previous reports comparing temporal clustering between single- and dual-task encoding conditions (Mundorf, Uitvlugt, & Healey, 2022; Healey, Long, & Kahana, 2019). No significant differences in FTA were observed between the encoding conditions, *t*(59) = .69, *p* = .5, *d* = .21. Aside from exhibiting greater temporal contiguity, single-task recall performance was similar to that observed under dual-task encoding.

### EEG Results

#### Subsequent Memory Effects

To examine the effects of dual-task performance and target detection on neural SMEs, we focused our analyses on a medial parietal region of interest (ROI; Figure 1a) where alpha and gamma band SMEs have been previously observed (Weidemann & Kahana, 2021; Sederberg et al., 2006). The ROI included 13 channels (Pz, P1, P2, P3, P4, CPz, CP1, CP2, CP3, CP4, P0z, P01, and P04). SMEs were computed using two steps: 1) separately averaging the observed power in each of the frequency bins over encoding trials for subsequently remembered and forgotten items, and 2) calculating the averaged remembered minus forgotten trial power. Table 1 reports numerical SMEs for each frequency band and Figure 2 illustrates the full time-frequency spectrograms for each encoding condition.

**Figure 2.**
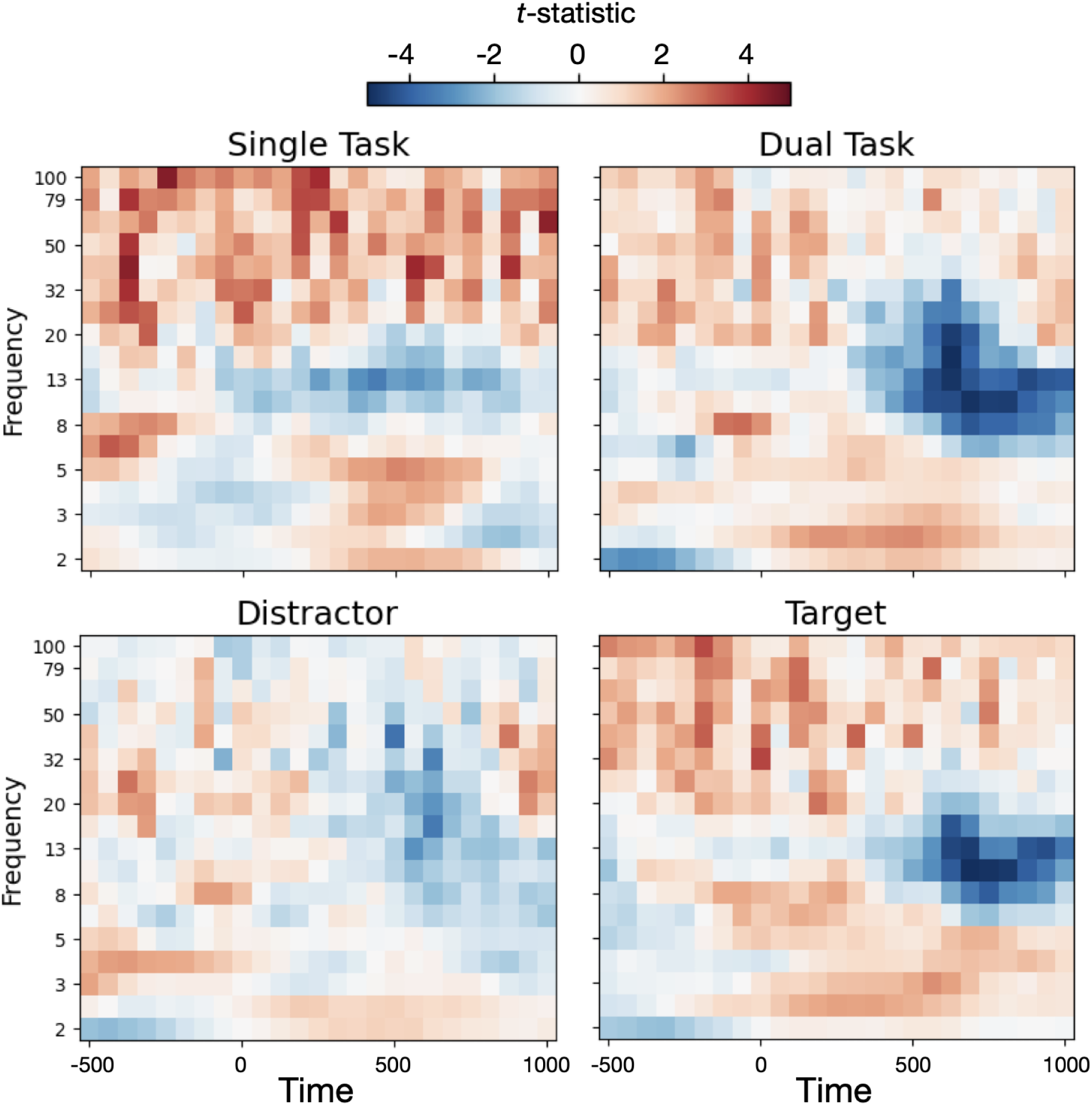
Spectrogram *t*-statistics for remembered vs. forgotten trials in the bilateral parietal ROI. Subsequent memory effects are plotted separately for dual-task and single-task (top) encoding conditions, and for the target and distractor trials within dual-task participants (bottom). All plots range from -500 ms to 1000 ms across trial onset.

**Table 1.** Across-participant *t*-statistic (*p value*) for Remembered vs. Forgotten Z-power spectral density within each frequency band.

| Freq. Band | Distractor | Target | Single Task | Dual-Task |
| --- | --- | --- | --- | --- |
| Theta (4-8 Hz) | -1.8 (.08) | .56 (.58) | 1.2 (.26) | -.27 (.79) |
| Alpha (8-12 Hz) | -1.9 (.07) | **-3.9 (< .001)** | **-2.26 (.04)** | **-3.8 (< .001)** |
| Beta (12-30 Hz) | -1.9 (.07) | -1.6 (.11) | .18 (.85) | **-2.9 (< .01)** |
| Low Gamma (30-50 Hz) | -1.9 (.07) | .93 (.36) | **3.8 (< .01)** | -.47 (.64) |
| High Gamma (50-100 Hz) | -1.2 (.23) | 1.4 (.16) | **4.0 (< .001)** | .53 (.60) |

Note: Means and CIs were computed from the across-participant averages. Bold font indicates the SME significantly differs from zero at $p < .05$ .

We first sought to determine whether previously observed SMEs replicated among participants in the single-task condition. To do this, for each participant we examined neural activity from word presentation within the alpha and high gamma frequencies. In accordance with prior work (Long et al., 2014; Ezzyat et al., 2017) we first calculated a t-statistic contrasting remembered vs. forgotten trials within each participant in each of the frequencies. This procedure accounts for across-trial variance in the magnitude of these effects to produce a subject-level SME at each frequency. The resulting t-statistics were then submitted to one-tailed across-participants t-tests. Among participants in the single-task condition, *t*-tests indicated a negative SME in the alpha band, Remembered (R) - Forgotten (F) *t*(20) = -2.2, *p* = .04 and a positive SME in the high gamma band, R - F *t*(20) = 4.0, *p* < .001. We therefore replicated the gamma and alpha band SMEs in the single task condition.

Of greater interest were the effects of dual-task performance on these SMEs. Prior work manipulating how the words themselves are processed suggests that increasing cognitive load should reduce gamma band SMEs (Long & Kahana, 2017; Broitman et al., *under review*). Consistent with this possibility, among dual-task participants a significant positive SME within the high gamma band was not observed, *t*(39) = .53, *p* = .60. The negative alpha band SME was replicated, R - F *t*(39) = -3.8, *p* < .001. To determine whether SMEs differed between the single- and dual-task conditions, we performed contrasts on the alpha and gamma MLM’s. While the alpha band SME did not significantly differ between single- and dual-task encoding conditions, z(80408) = .62, p = .53, the high gamma SME was significantly more positive in the single-task condition than in the dual-task condition, z(80420) = 2.7, p < .01. This result indicates that associations between high gamma activity and memory encoding are disrupted under divided attention.

To determine whether the differences observed between the single- and dual-task high gamma SMEs were specific to target detection or distractor rejection, we separately examined data from target and distractor trials among dual-task participants by performing a contrasts analysis on the Gamma MLM. Contrasts revealed a significant interaction of cue condition and subsequent memory, *z*(1, 80408) = 3.5, p < .001, distractor R - F *d = -*.01, target R - F *d =* .04, such that the high gamma SME was more positive during target trials than distractor trials. The effect of dual-task encoding on gamma SMEs therefore appears to be stronger on distractor trials.

#### Changes in Subsequent Memory Effects Over Time

A central goal of this study was to characterize the time course of the neural SMEs. We therefore asked whether the spectral SMEs occurred pre- or post-stimulus by dividing the data from each trial into 3 time bins: a 500 ms pretrial period (*pre-trial*), prior to the onset of the encoding word and detection cue; the 500 ms period following the trial (*early trial*); and the period between 500 and 1000 ms following trial onset (*late trial*).

#### Alpha Band

Because alpha band activity is associated with attentional orienting and successful encoding, we next examined how alpha power changes over time. Alpha band power is visualized by time bin, trial condition, and subsequent memory in Figure 3a. For a visualization, Figure 3b plots alpha power across cue conditions, subsequent memory, and time bins in the *a priori* parietal ROI.

**Figure 3.**
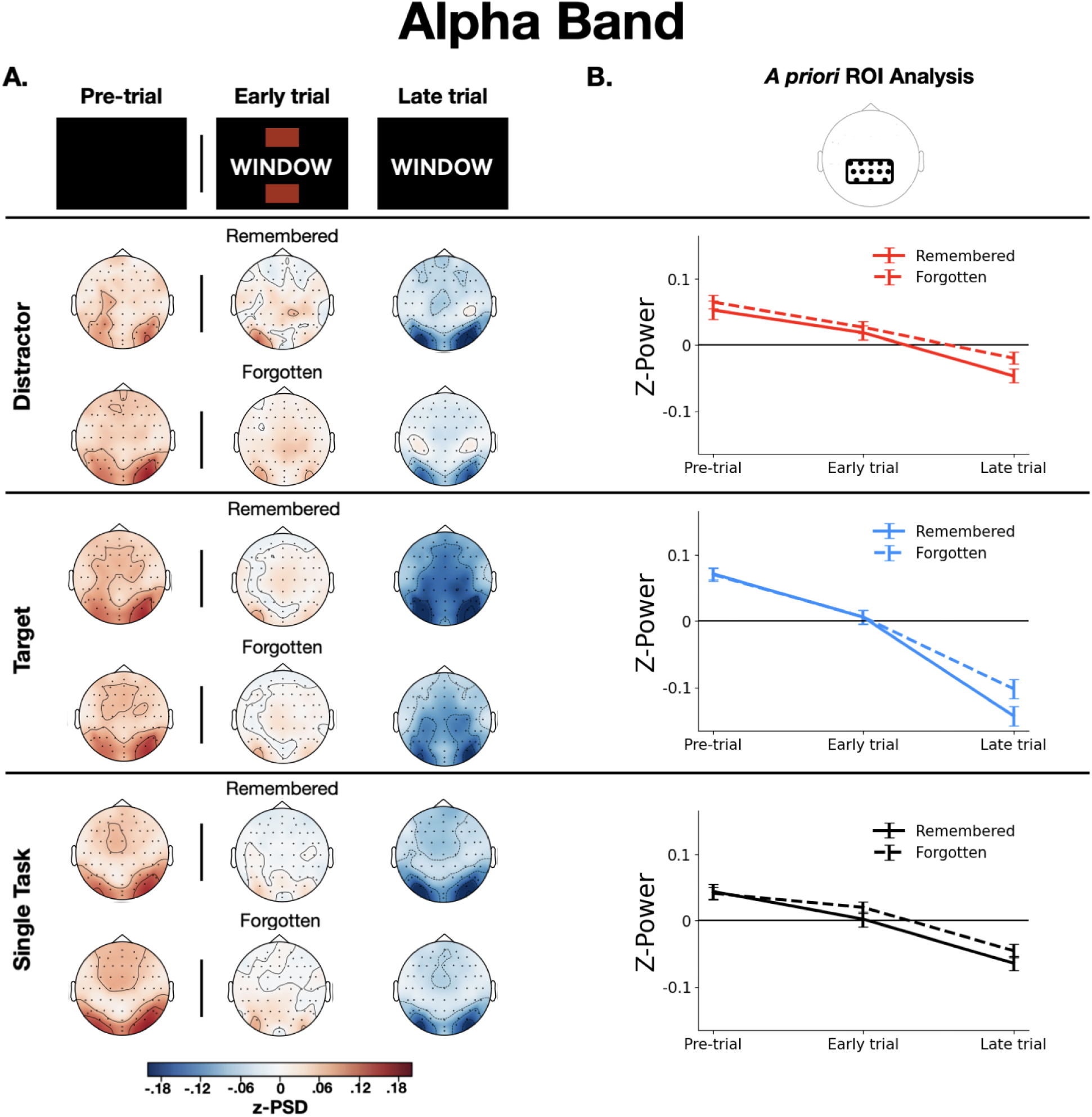
**A)** Topographic plots of alpha band activity across trial time bins by trial condition (distractor, target, and single task) and subsequent memory (remembered and forgotten). **B)** Line plots of alpha activity within the *a priori* parietal ROI. Error bars reflect the standard error of the mean computed across participants.

Prior reports suggest that attention-related alpha suppression effects are frequently observed *after* stimulus presentation (Klimesch et al., 1998; de Graaf & Thompson, 2020; He et al., 2021), and that alpha band SMEs are typically observed more than 1000 ms following trial onset (Noh et al., 2014; Fellner et al., 2019). This suggests that any effects of cue condition on SMEs would be likely to emerge after stimulus onset. We therefore investigated the interaction of cue type and subsequent memory in the two post-stimulus time bins using tailored contrasts with the Alpha MLM. Among dual-task participants, a main effect of cue condition was observed during the early trial bin, *z*(2, 80408) = 10.6, p < .0001, target-distractor *d* = -.25. Furthermore, when alpha activity was examined specifically in the late trial bin, the SME for target-paired words was more negative than that of Distractor-paired words, interaction of cue condition and subsequent memory, *z*(2, 80408) = 3.2, p = .001, distractor R - F *d = -*.24, target R -F *d = -*.33. The results suggest that lower levels of alpha band activity following target detection, relative to distractor rejection, are related to successful memory encoding.

The time course of alpha band activity changes in each condition was further characterized by applying additional contrasts to the Alpha MLM. Across all dual-task trial conditions (remembered vs. forgotten, target vs. distractor), stimulus onset resulted in a drop in alpha band activity, F(2, 115) = 72.5, p < .0001. However, this drop was greater for items presented with a target, interaction of time and cue condition, *z*(2, 80408) = 15.1, p < .0001, distractor late trial - pretrial *d = -*.29, target late trial - pretrial *d = -*.63, as well as for words that were subsequently remembered, interaction of time and subsequent memory, *z*(2, 80408) = 4.5, p < .0001, R late trial - pretrial *d = -*.57, F late trial - pretrial *d = -*.48. Finally, a three-way interaction of time, trial type, and subsequent memory indicated that target-paired words had a greater alpha band SME toward the end of the trial, *F*(4, 52063) = 4.1, *p* = .01.

Figure 3 suggests that the drop in power following stimulus presentation in the single-task condition was similar to that observed during distractor trials in the dual-task condition. To evaluate this effect quantitatively, we separately compared single-task alpha band activity with that of target and distractor trials. Changes in alpha band activity over time during single-task trials did not significantly differ from distractor trials, *z*(2, 80408) = .17, *p* = .87, distractor late trial - pretrial *d’s* > -.29, single task late trial - pretrial *d* = -.28, but did differ from target trials, *z*(2, 80408) = 5.2, *p* < .0001, target late trial - pretrial *d* = -.62. Because alpha band activity decreases when orienting to external information (e.g., Klimesch et al., 1998), this finding is consistent with target detection eliciting a reactive response that facilitates stimulus processing and encoding.

#### High Gamma Band

Another set of analyses examined whether high gamma band power is modulated by divided attention and target detection at different times relative to the onset of the stimulus. Averaged high gamma band power is visualized by time bin, cue type, and subsequent memory in Figure 4b.

**Figure 4.**
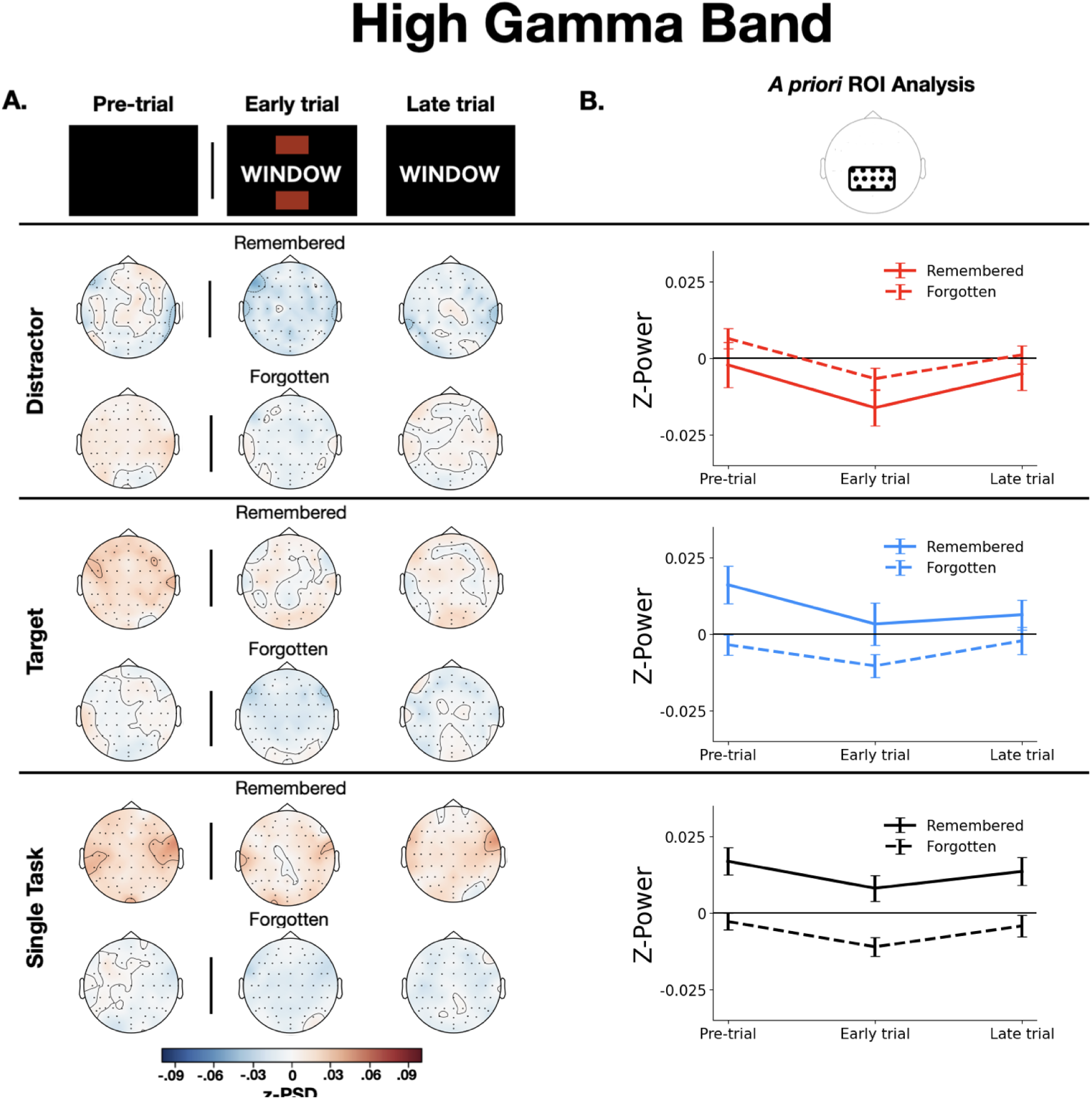
**A)** Topographic plots of high gamma band activity across trial time bins by trial condition (distractor, target, and single task) and subsequent memory (remembered and forgotten). **B)** Line plots of high gamma activity within the *a priori* parietal ROI. Error bars reflect the standard error of the mean computed across participants.

Previous studies of gamma band SMEs suggest that they may be detectable prior to stimulus onset (Noh et al., 2014), consistent with an association with the maintenance of task set. To evaluate whether pre-stimulus states contribute to SMEs and the effects of target detection on encoding, we examined dual-task high gamma power specifically in the pre-trial time bin. Though there was no overall pre-trial SME, there was an interaction of cue type and subsequent memory, *z*(2, 80420) = 2.3, p = .026, distractor R - F *d* = -.03, target R - F *d* = .06. When all time bins were considered, an interaction of cue type and subsequent memory indicated that the positive high gamma band SME for target trials significantly differed from the negative SME for Distractor trials, *z*(2, 80420) = 3.5, *p* < .001, distractor R - F *d* = -.02, target R - F *d* = .06. In addition to these interactions, the Gamma MLM revealed main effects of recall status, indicating a positive SME across all trial types, *F*(1,59) = 5.1, *p* = .03. A main effect of time bin reflected a dip in gamma power immediately following stimulus onset across all trial types (Figure 4), *F*(2, 80442) = 7.9, p < .001. These results show that gamma SMEs were present across time bins and significantly differed for target and distractor trials, even before the onset of the cue.

To determine whether the effects of target detection on memory varied across trial types, we conducted separate statistical contrasts comparing high gamma activity in the single-task condition to the target- and distractor conditions across the full trial duration. While the high gamma band SME was significantly greater for the single-task than for distractor trials, significant interaction of trial type and subsequent memory, *z*(2, 80420) = 2.5, p = .01, distractor R - F *d* = -.02, single task R - F *d* = .06, it did not significantly differ from that on target trials, nonsignificant interaction of trial type and subsequent memory *z*(2, 80420) = .46, *p* = .64. These results indicate that single-task high gamma activity was similar to that of the target trials across the full trial duration.

#### Post-hoc clustering analysis

A visual inspection of the target and distractor topographic plots in Figures 3 and 4 indicated that interactions between cue type and subsequent memory may differ among channels that were not included in the *a priori* ROI. To investigate whether this was the case, we conducted a *post hoc* clustering analysis with data from the dual-task encoding condition. Using MNE’s permutation clustering function, we first identified groups of adjacent electrodes that demonstrated SMEs in a given frequency band beyond a threshold of p < .001 (*clusters*), collapsing across cue conditions. The observed t-statistics of the identified clusters were then compared to 1024 randomly permuted clusters created by shuffling the participant and condition (remembered/forgotten) labels for each data point. In the alpha band, the permutation test identified two significant clusters of electrodes in the occipital and left frontal areas. Results from these clusters were largely consistent with those from the *a priori* parietal ROI: both clusters demonstrated an alpha band SME, an effect of time bin, a time x cue condition interaction, and a time x subsequent memory interaction (all p’s < .003). The results suggest that the interactions of cue type and subsequent memory are not limited to the medial parietal channels included in our *a priori* ROI, but also may be present in occipital and frontal scalp sites.The same analyses were performed with high gamma power as the dependent variable, but no significant clusters were observed.

## Discussion

The present study investigated whether the neural correlates of memory encoding reflect variability in attention over time within the context of the attentional boost effect. Although SMEs were evident under single-task encoding conditions, they were significantly modulated by the need to perform the secondary target detection task. Our findings further revealed that target detection itself reduced alpha band power, and this reduction was associated with successful encoding. We also observed high gamma band SMEs during target and single-task trials, but not during distractor trials. Importantly, differences in gamma band power were apparent prior to stimulus presentation, which could reflect a role for pre-trial attentional states in memory encoding and in the detection of behaviorally relevant stimuli such as targets. Taken together, the results point to multiple mechanisms by which momentary changes in attention can influence episodic encoding.

### Divided attention alters gamma SMEs

Data from the single-task encoding condition replicated prior findings that increases in high gamma and decreases in alpha power predict subsequent memory (e.g., Sederberg et al., 2003; Long et al., 2014; Fellner et al., 2019). However, the high gamma SME was not observed under divided attention, which extends prior work (Long & Kahana, 2017; Broitman et al., *under review*) to dual-task situations that require the processing of multiple, simultaneously presented stimulus streams. Because gamma oscillations appear to be important for maintaining task-oriented states (Clayton et al., 2015; Randazo et al., under review), smaller gamma SMEs under divided attention could reflect the disruption of those states when two tasks are performed at once.

Importantly, the effects of dual-task encoding on gamma SMEs differed across the target and distractor conditions. Specifically, as in the single-task condition, high gamma activity preceded encoding success during target trials. This was not the case during distractor trials, suggesting that the attentional states that produce gamma SMEs may be beneficial for memory encoding during target presentations but not during distractor presentations. Further, although responding to targets requires greater attention than rejecting distractors (Duncan, 1980; Raymond et al., 1992), the current results suggest that target detection does not disrupt cognitive states that facilitate encoding.

The lack of a high gamma SME for distractor trials is surprising. It suggests that successful encoding when a distractor appears relies on different processes than those engaged under single task or target conditions. Behaviorally, most prior evidence suggests that distractor rejection produces little interference in the ability to encode coinciding items above and beyond the effects of dual-task interference (Lee, 2023; Moyal et al., 2022; Mulligan, et al., 2014; Swallow & Jiang, 2014), though small effects have been reported (Meng, et al., 2019). However, rejecting an item as a distractor interferes with memory for the item itself (Chiu & Egner, 2015). It therefore may be possible that the lack of a gamma SME on distractor trials reflects the effects of interference from distractor processing. However, because no dual-task baseline condition was included in this study, we cannot disentangle the effects of interference from performing two tasks at once from the effects of processes engaged when a distractor appears (c.f., Lee, 2023; Moyal, et al., 2022; Swallow & Jiang, 2014). Additional research is needed to further characterize the interaction between pre-stimulus attentional states and distractor processing.

Considered with evidence that gamma activity reflects the ability to maintain task readiness over an extended duration (Clayton et al., 2015), our results suggest that preexisting attentional states may influence an individual’s ability to select and attend to behaviorally relevant events when they occur. Target detection therefore might allow individuals to take advantage of the same conditions that facilitate encoding under full attention. Meanwhile, the absence of such effects prior to distractor presentation suggests that being in a “readiness to encode” state offers little advantage when one must process and reject a distractor.

### How does target detection boost memory?

A central goal of this study was to investigate the effects of target detection on EEG spectral features that have previously been associated with successful encoding. The results may lend insight into the competing accounts of whether target detection elicits cognitive states consistent with single task encoding, or whether it elicits a mechanism that temporarily boosts perceptual processing and encoding (temporal selection).

If target detection alleviates dual-task interference, then we should expect to see patterns that are consistent with single-task SMEs following target detection (i.e., post-target increases in gamma power and decreases in alpha power). Although targets elicited decreases in alpha power, we found no evidence of post-stimulus changes in high gamma activity related to subsequent memory. Instead, we found that high gamma power predicts memory encoding prior to the onset of target and single-task trials, which is consistent with findings of pre- and early-trial SMEs in the gamma band (Noh et al., 2014; Fellman et al., 2019). This result is consistent with work that suggests that the effects of LC-mediated orienting responses are modulated by prior attentional states (Mather et al., 2016, Swallow et al., 2022).

Our finding that target detection reduced alpha power, and that larger reductions were associated with more successful encoding, may provide evidence for temporal selective attention as proposed by the DTI model (Swallow & Jiang, 2013). The results align with previous research that has linked alpha oscillations with externally oriented attention, phasic LC activity, and the detection of behaviorally relevant stimuli (Compton et al., 2021; de Graaf & Thompson, 2020; He et al., 2021). The effects reported here may therefore reflect the prioritization of information presented concurrently with targets, and are consistent with prior pupillometric and fMRI work (Moyal, et al., 2022; Swallow, Jiang & Riley, 2019; Yebra, et al., 2019).

### Study limitations

Several components of the current experiment may have limited our ability to fully capture the effects of target detection on encoding. First, the number of participants, encoding trials, and scalp electrodes were small compared with some previous EEG studies of episodic memory (e.g., Long et al., 2014; Sederberg et al., 2006). Additionally, the encoding trial durations were brief compared to most other studies of neural subsequent memory effects (e.g., Fellner et al., 2019; Sederberg et al., 2003; Long et al., 2014). Although our results demonstrate that alpha and high gamma SMEs are robust and highly replicable, it is possible that more subtle effects would have emerged with greater power.

Additionally, relative to single-task, list length was shorter in the dual-task condition to better equate recall rates across conditions. However, differences in list length seem an unlikely source of our findings for two reasons. First, the effects of divided attention and target detection on memory performance were consistent across serial positions. Consequently, early and late list positions contributed similarly to the SMEs across encoding conditions. Second, the standard spectral SME, higher gamma and lower alpha power, is most prominent in early list positions (Sederberg et al., 2006), before any effects of list-length could occur. Though these findings are reassuring, a systematic investigation of the effects of target detection on spectral EEG components across lists and list positions is needed to elucidate the influence of attention on memory encoding at multiple time scales.

Finally, target trials required a button press while distractor and single-task trials did not. This raises the possibility that motor processing contributed to our findings. Several factors that arbitrate against the possibility that our results can be attributed to the button press itself. First, novel effects reported here either occurred prior to trial onset and before the appearance of a target or distractor (in the case of pre-stimulus gamma SMEs) or involved within condition SMEs that matched on the presence or absence of a button press (in the case of post-stimulus alpha SMEs). Second, the effects of target detection on both high gamma and alpha power were spread across multiple scalp sites that extended well beyond regions likely to show sensorimotor effects (Dieber, et al., 2012; Papausek & Schulter, 1999), and included a bilateral occipital cluster (Figures 3 and 4). Finally, although producing the attentional boost effect does not require that participants perform a physical action (Swallow & Jiang, 2012; Mulligan et al., 2016; Swallow et al., 2019), any residual effect of motor processes that is not addressed by these points (pre-stimulus effects, effects when conditions are matched by button press presence or absence, effects observed diffusely and in occipital regions) suggests a greater connection between motor processing and memory than a simple effect generated by the movement itself (see, e.g., Yebra, et al., 2019). Future research should further investigate this possibility.

### Directions for future research

We specifically focussed on free recall SMEs in the alpha and high gamma bands because both effects can be reliably detected in scalp EEG, and because those frequency bands have distinct associations with changes in attention. However, other kinds of oscillations have also been shown to interact with both attention and memory encoding. Delta band activity (2-4 Hz) positively correlates with encoding success and may be sensitive to attentional fluctuations across list positions (Sederberg et al., 2006). Theta band oscillations (4-8 Hz) have been observed to have both positive (Klimesch et al., 1997; Osipova et al., 2006) and negative (Guderian et al., 2009; Long et al., 2014) associations with encoding success in scalp EEG studies. Frontal theta rhythms have also been observed to increase under cognitive control (Cavanagh & Frank, 2014), and may be sensitive to shifts in attention over spatial locations (Fiebelkorn & Castner, 2019). Thus, theta oscillations may reflect the operation of multiple neurocognitive mechanisms involved in memory formation (for a review, see Herweg et al., 2020). Finally, EEG data also contain aperiodic signals that are modulated by attention and task demands (Deodato & Melcher, 2024). Future studies may benefit from investigating a broader range of oscillatory and aperiodic activity in order to gain a more comprehensive understanding of the mechanisms involved in attention and memory interactions.

Future studies may also want to examine the specific role of distractor rejection and dual-task interference in high gamma SMEs. Adding a no-cue condition under dual-task encoding could help to clarify whether target detection alleviates dual-task interference, or distractor rejection specifically reduces the role of gamma activity in encoding success. Finally, the causal role of brain activity related to encoding success in memory improvements is a topic of discussion in the current literature (see Halpern et al., 2023). Although there is some evidence that this brain activity can directly increase episodic encoding in intracranial EEG (Ezzyat et al., 2017; 2018), this has yet to be demonstrated using scalp EEG. Future studies may benefit from applying a “closed loop” approach to stimulus presentation, in which neural data are continuously read through a computer algorithm that predicts whether items presented in a given moment will be successfully encoded.

### Conclusions

The ability to sustain attention to a task, divide attention across tasks, and orient to behaviorally relevant stimuli can each impact memory encoding. The current study used EEG to better characterize the impact of each of these factors on neural subsequent memory effects in the context of the attentional boost effect. We found evidence that pre-stimulus task engagement and target detection modulate the relationship between successful encoding and gamma band and alpha band activity. The results suggest that neural SMEs reflect multiple sources of variability in attention over time.

## Acknowledgements

The authors would like to thank Amy Krosch for allowing us to use her EEG facilities and Sanweda Maghabin, Jackie Kim, Sahib Kaila, for assistance with data collection on this project. Funding for this project was provided by the Cornell University College of Arts & Sciences.

## Data Availability

Data are available upon reasonable request and with proper approval from relevant research and ethics entities. Requests should be directed to Khena Swallow. Several key EEG preprocessing and data analysis scripts are available for download at https://github.com/awb99cu/EEG_Free_Recall_Scripts.

